# Programmed inhibition of an innate immune receptor via *de novo* designed transmembrane proteins

**DOI:** 10.1101/2025.05.31.657109

**Authors:** Colleen A. Maillie, Minghao Zhang, Nadia Gosiet, George Goldenfeld, Andrew B. Ward, Marco Mravic

## Abstract

Transmembrane domains of immune complexes transmit precise signals across lipid bilayers. Probing their interactions has the potential to yield mechanistic insights relevant to therapeutic design. However, our capability to generate molecules directed to bind lipid-embedded sites is limited. Here, we demonstrate a computational strategy to design polypeptides targeting Toll-like receptor 4 (TLR4), a mediator of inflammatory signaling, directly within membranes. TLR4 poses a formidable molecular recognition challenge, as its transmembrane domain is largely apolar, lacks a defined sequence motif, and exhibits an underdetermined structure–function relationship. One synthetic protein binds TLR4’s transmembrane domain and antagonizes NFκB signaling in human cells, proving that precise transmembrane domain interactions are essential for cross-membrane conformational coupling. This work refines design principles for encoding stable interactions in cellular membranes and expands the range of lipid-embedded mechanisms accessible to probe with computationally derived molecules.

## Introduction

Immune receptors comprise membrane-embedded signaling hubs to detect extracellular stimuli and initiate intracellular signal transduction. These immune communication sites often require precise molecular interactions between transmembrane (TM) domains to modulate and tune diverse signaling pathways across the lipid bilayer^1,2^. Despite their functional importance, receptor membrane-spanning regions remain poorly understood, and conventional water-soluble molecular tools have limited ability to manipulate these regions due to their embedding within the lipid bilayer. Targeting proteins from within the membrane itself offers an innovative strategy for modulating receptor activity^3–7^. This approach remains largely untapped, but is becoming increasingly accessible through computational protein design^8–12^.

In this study, we sought to advance chemical methods of targeting TM domains through the process of binding an immune receptor, Toll-like receptor 4 (TLR4). TLR4 is a self-versus non-self-sensing receptor best known for its role in detecting bacterial pathogens and triggering an innate immune response through pro-inflammatory pathways^13–15^. Beyond bacterial infection, TLR4 signaling is critically implicated in diseases such as lupus, sepsis, allergies, and colitis, where receptor dysregulation presents a key target for intervention^16–19^. Additionally, therapeutic strategies for infectious diseases, cancers, and autoimmune disorders leverage TLR4 modulation^18,20–23^.

Despite the central role for TLR4 in immunity and disease, major gaps remain in our understanding of the structure–function relationships that govern its activation mechanism. Current models are largely focused on the conformations of the ectodomains and imply long-range cross-membrane geometric coupling^24^. X-ray crystallography and cryo-electron microscopy studies show lipopolysaccharide (LPS) agonist-bound extracellular domains dimerize alongside co-receptor myeloid differentiation factor 2 (MD2) while apo or antagonist-bound states adopt monomeric ectodomain arrangements^25–28^. However, FRET-based microscopy in live, unstimulated cells revealed that nearly half of the TLR4 population may exist as preformed dimers^29^. Such data challenges the simple ligand-induced dimerization model and supports a more complex mechanism of preassembled complexes activated by conformational change. How the membrane-spanning region of TLR4 contributes to transmitting ectodomain structure into intracellular signaling efficacy and bias is poorly defined, although the resulting TM geometry appears impactful. Truncation of TLR4 extracellular domains leads to constitutively active receptors^30^. Similarly, uncoupling the ecto- and TM domains with flexible linkers results in constitutive activation, bypassing ligand-dependent activation^31^. Moreover, isolated TM regions from the TLR family members, including TLR4, have been shown to homo- or hetero-dimerize in bacterial membranes^32^. Mimic TM peptides of TLR2 function as dominant negative inhibitors of signaling, likely via these TM interactions^33,34^. Solution NMR has provided insight into the oligomerization and extended helical architectures of TM and juxtamembrane (JM) regions^35,36^. Despite this evidence, the relevance of TM domain self-interactions and conformations in full-length TLR4 during signaling within mammalian cells remains unclear and mostly untested.

Although TLR4 plays a central role in inflammatory signaling, it is an underexploited therapeutic target. While vaccine adjuvant strategies successfully leverage TLR4 agonism, no FDA-approved small molecule antagonists currently exist to address TLR4’s role in a myriad of inflammatory disorders^20,37–39^. Chemical matter engaging TLR4 has been largely restricted to lipid endotoxin scaffolds and derivatives^40^. Likewise, mechanistic and structural complexities of the receptor-ligand interactions in signaling have limited effective rational drug development. TLR4 activation is governed by fine-tuned molecular features, exemplified by the remarkable stereoselectivity of the synthetic agonist Neoseptin-3; a single chiral center inversion can fully abolish its biological activity^28,41^. Similarly, single-point chiral modification of lipid A analog CRX-547 can bias downstream signaling from MyD88-to TRIF-dominant pathways^42^. Clarifying key aspects of its ligand-induced structural mechanism could significantly improve our abilities to chemically modulate TLR4 for therapeutic benefit.

To expand the repertoire of molecular probes available and to dissect TM determinants signaling, we computationally designed synthetic polypeptides aimed at binding the membrane-embedded surface of TLR4. TLR4 poses a molecular recognition challenge reaching beyond the current cutting-edge for targeting in lipid bilayers^9,10,12^. Like the TM spans of most TLR family members, its sequence is largely apolar and lacks patterns or conserved motifs well-recognized amongst membrane proteins characterized to date^43^. As well, the helix-helix interface conformation and key residues mediating TM domain dimerization and functional signaling in mammalian cells have been proposed from modeling and *in vitro* experiments but have not been confirmed experimentally^35,44^. Despite lacking precise structural information, we hypothesized that the TM helix of TLR4 should be a tractable site for synthetic targeting, given it encodes specific interactions required for assembly and activation. Likewise, if TM domain interactions are involved in signaling, then transmembrane peptides competitively binding with the native dimerization would inhibition receptor signaling. We developed a computational design framework based on membrane-specific chemical principles to generate sequences targeting TLR4’s TM span. The lead resulting polypeptide forms a stable TM complex both in cells and *in vitro*, leading to significant dampening of TLR4-mediated NFκB signaling in mammalian cells. Our results determine that endogenous TM conformations regulate TLR4 activation in cells, thus highlight a new strategy for dissecting its signaling mechanism and re-programming immune signaling via lipid-embedded molecules generated on-demand. This work advances the technological capabilities for molecular targeting in membranes with minimal prior structural information, broadening the scope of surface receptors that rely on cryptic transmembrane signaling mechanisms accessible to probe for therapeutic potential, like TLRs.

## RESULTS

### TLR4 transmembrane domain self-interaction and its role in signaling

We hypothesized that if TLR4 requires transmembrane homodimerization for signaling, then protein fragments spanning TLR4’s TMJM segment could competitively engage these interactions, disrupt an active dimer complex, and inhibit NF-κB signaling downstream of LPS stimulation (Figure 1A). First, to test whether stretches of TLR4’s membrane-spanning domains self-interact in lipid bilayers of mammalian cells, we implemented a variant of the split enzyme complementation NanoBit assay we recently validated for membrane-embedded interactions^45^. Protein constructs including fragments of TLR4 spanning the TM domain 20-aa central stretch “TLR4-TM” or extended to 35-aa including the juxtamembrane (JM) region “TLR4-TMJM” were fused at the C-terminus with LgBit and SmBit enzyme fragments and co-expressed together, or expressed with negative-control, non-interacting LgBit-fused TM proteins (CD8α, eVgL^46^) (Fig. 1B). These constructs express well with plasma membrane localization in HEK293T cells (Supplemental Figure 1A,C-D). In this assay, a primary luminescence event indicates the protein-protein interaction (PPI) propensity. Then, after introducing the self-complementing HiBit peptide to displace smBit, a secondary luminescence event is read to quantify the total LgBit-fused protein present, then used to normalize the observed PPI luminescence by each protein’s underlying expression level. This expression-normalized PPI signal corrects for expression level differences of enzyme components to better represents protein interaction propensity. Homotypic interactions upon co-expression of pairs of TLR4-TM or TLR4-TMJM SmBit and LgBit constructs result in expression-normalized luminescence that is far greater than when SmBit-tagged TLR4-TM or TLR4-TMJM proteins are co-expressed with non-interacting proteins eVgL-LgBit (>100-fold change in signal, p < 0.005, n=3) and CD8α-LgBit (2.2 ± 0.3 fold, p < 0.005; 1.8 ± 0.3 fold, p < 0.05, respectively) (Figure 1B, Supplemental Figure 1B,E). Thus, at moderate expression levels within this experiment, TLR4’s isolated TM spans inherently self-associate in mammalian cells. Given this robust self-assembly, we expect that TLR4’s parallel lipid-embedded interaction is physiologically relevant and may be a favorable driving force for ligand-independent receptor dimerization, thus must be opposed by stronger forces from other receptor domains to prevent aberrant signaling (i.e. ectodomain steric exclusion or conformations incompatible with the TM geometry). Furthermore, we provide evidence that this TM domain interaction and its associated energetics contribute significantly to TLR4’s structural mechanism, both during ligand-induced ectodomain rearrangement and in mediating long-range interdomain coupling that links ligand identity to downstream signaling outcomes.

**Figure 1.**
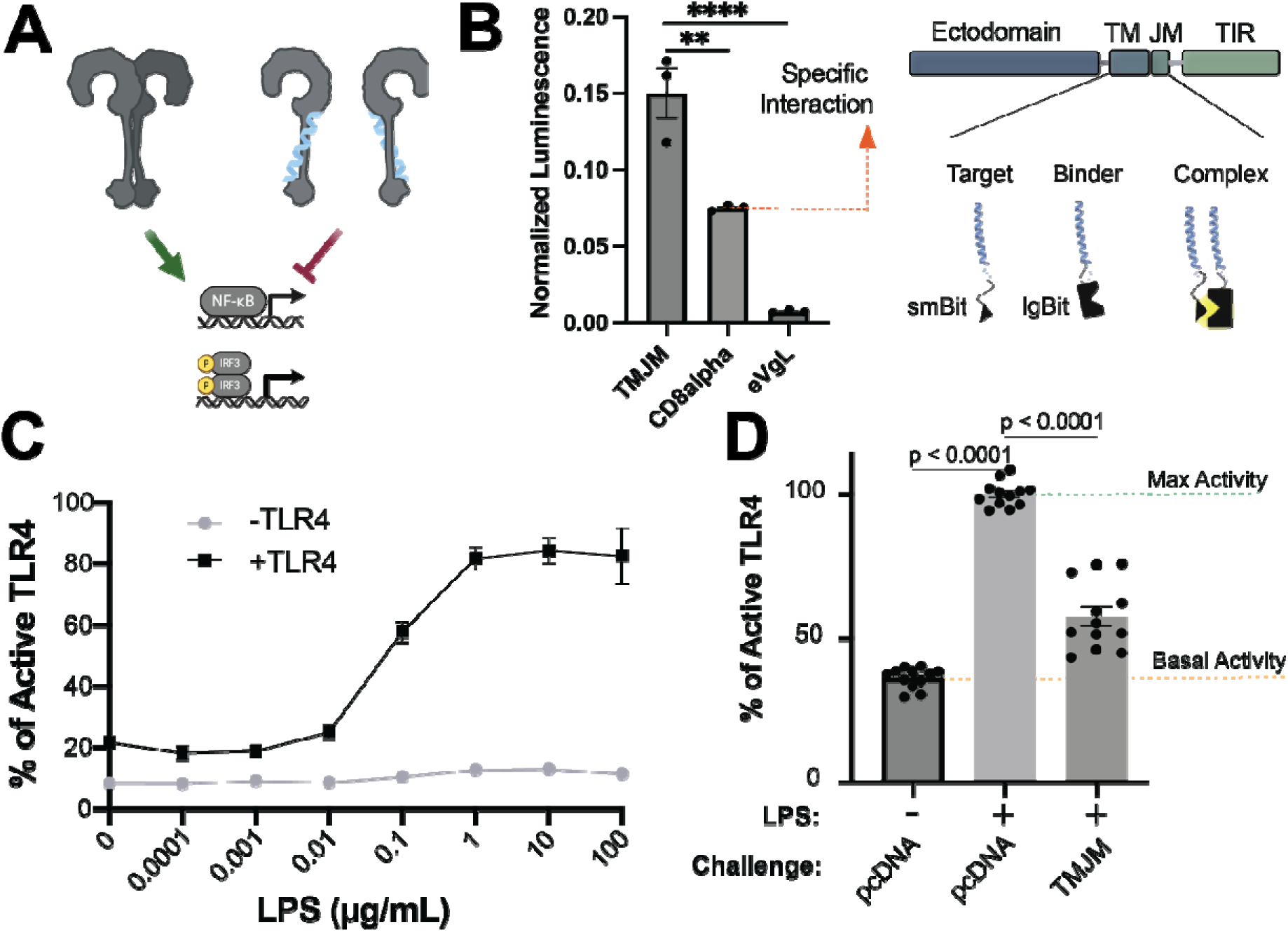
TLR4 TM domains self-interact and can antagonize LPS-induced signaling. **A)** Proposed model for TLR4 signaling uses native transmembrane (TM) domain interactions in the complete active receptor complex for transmitting signals across bilayers. Exogenous TM peptides targeting TLR4 in membranes should disrupt downstream receptor signaling. **B)** Left, expression-normalized NanoBiT luminescence co-expressing isolated TLR4-TMJM-SmBit with TLR4-TMJM-LgBit or compared to co-expressing non-interacting controls (CD8α-LgBit, eVgL-LgBit) in HEK293T cells. Dots are technical triplicate in 1 biological replicate representative of n=3 total independent trials. Asterisks denote two-tailed p-value of significance by student’s t-test (**, 0.05; ****, <0.0001). Error bars, standard error of the mean (SEM). Orange dash line (CD8α-LgBit) represents normalized PPI signaling of non-specific complementation baseline. Right, TLR4 domain architecture and cartoon schematics of TM NanoBiT constructs. **C)** LPS dose-response in HEK293T/MD2/CD14 TLR4-null NF-κB-SEAP reporter cells upon transfecting TLR4-eGFP or empty vector, normalizing % of active TLR4 by maximal response at 100 µg/mL LPS stimulation. Error bars, SEM. Representative of n=3 biological replicates. **D)** NF-κB-SEAP reporter signal upon co-transfection with TLR4 alongside either empty pcDNA vector or TLR4-TMJM fragment protein-containing plasmid and stimulation with 10 µg/mL LPS. Dots are technical triplicates from *n* = 3 biological replicates. Error bars, SEM. P-value, significance from two-tailed student’s t-test. Percent of active TLR4 was normalized to TLR4 activity stimulated with 10 µg/mL LPS.

We next evaluated the capacity of inter-molecular interactions from the TLR4-TMJM polypeptide to modulate signaling of the full-length TLR4 upon co-expression. We monitored agonist-stimulated TLR4 signaling in a NF-κB-inducible secreted embryonic alkaline phosphatase (SEAP) colorimetric reporter HEK293 cell line wherein endogenous TLR4 is absent and essential TLR4 co-receptors MD2 and CD14 are stably transduced. TLR4 is introduced by transient transfection with a C-terminal fluorescent fusion protein (eGFP), which does not perturb signaling^47^ and allows monitoring of TLR4 expression. When TLR4-eGFP is transfected alongside empty pcDNA carrier plasmid and activated with titration of LPS, the NF-κB-SEAP reporter is elevated by 63.7±3.7% (n=3) above the basal signal level at 10 μg/mL agonist (Figure 1C,D). When the TLR4-TMJM polypeptide (LgBit fusion construct, as in Fig 1B) is co-transfected with TLR4-eGFP, we observed a 41.2±11.7% reduction in this endpoint NF-κB-SEAP reporter signaling at saturating LPS treatment (cellular response to LPS diminished to 22.5_±_12.8% (n=3); Figure 1D). Thus, challenge with TLR4’s TMJM fragment impairs maximum receptor signaling efficacy while the apparent EC_50_ for its ligand remains unaltered, consistent with inhibition mediated by the TM domain interaction and a dominant-negative effect. These findings reveal homotypic interactions that occur between TLR4’s TMJM region are influential to the full-length receptor’s cross-membrane conformational coupling. As such, we established that molecules intercepting the TM domain can be a viable mechanism to probe TLR signaling.

### Computational Design of TLR4 Targeting Peptides

Building on this mechanistic insight, we next explored whether TLR4 signaling could be modulated by synthetic polypeptides computationally designed to target its transmembrane region. TLR4’s TM span represents a challenge in lipid-embedded *de novo* molecular recognition, given its predominantly apolar sequence and lack of any sequence motifs known to encode TM dimerization. Design efforts to date has relied on recognizable sequence patterns of small or polar residues (Small-X_3_-Small, Small-X_6_-Small): as geometric templates during modeling and leveraging their known interaction propensities in lipid^8,9,48–50^. Likewise, neither the interaction geometry nor key residues mediating TLR4’s TM dimerization (i.e. those desirable to target) are fully clear or consistent when comparing results from modeling, co-evolutionary signatures, or previous NMR experiments (Supplemental Figure 2A). Thus previous design of TM proteins had utilized well-known sequence-structure relationships to inform binding interactions. By contrast, targeting TLR4 tackles the challenge of encoding a new stable complex in lipid with minimal existing structural information and driven mostly by complementary apolar packing.

Given TLR4’s TMJM region’s naturally evolved propensity for self-interaction, we posited that this molecular surface should be amendable to synthetic recognition and binding within lipid bilayers. Our approach to model TM helix interaction geometry was inspired by previous workflows used to engineer transmembrane (TM) domain mimics of BclxL proteins with improved affinity based on *ab initio* predicted homodimeric conformations^10^. Viable interaction geometries of the TLR4-TMJM homodimer complex were modeled by two orthogonal methods, AlphaFold2^51^ multimer or TMHop^52^, a Rosetta-based implicit membrane fold-and-dock algorithm. Amongst the molecular models sampled, there was no consensus or preferentially scored TM helix interface, underscoring the difficulty in identifying the most energetically favorable or physiologically relevant TM conformations this sequence (and those of mostly apolar composition). Therefore, we manually selected 3 predicted models as starting template backbone structures for computational design based on the rationale that those homo-dimer geometries pass common structural quality metrics (high interfacial interface surface area, high potential contact density). As well, these 3 unique binding modes were chosen since they engage different regions of TLR4’s TM helix (Figure 2A), thus cover distinct sets of residues possibly mediating its interface – spanning those co-evolving residues and those perturbed in TLR4 TM NMR oligomerization studies^35^ (Supplemental Figure 2A). Thus, this approach presents multiple opportunities to engage sites on TLR4’s TM region with functional consequence and provide sequence diversity.

**Figure 2.**
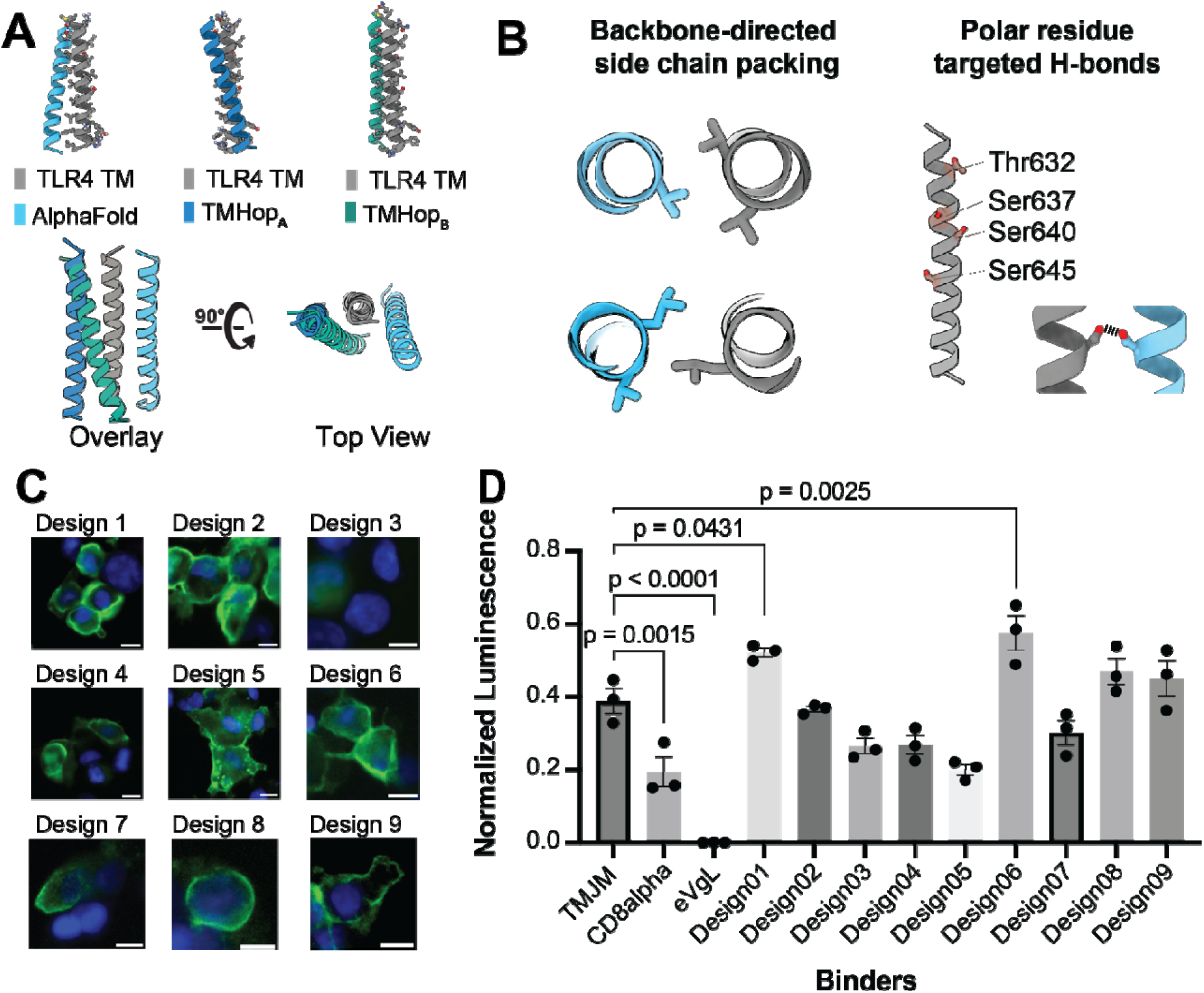
Computational design of *de novo* transmembrane polypeptides to target TLR4. **A)** Starting backbone models for anti-TLR4 TM-targeting designs were generated fromAlphaFold2 or TMHop TLR4 TM homodimeric predictions. **B)** *D*esign principles and molecular features optimized. Left, backbone-directed sidechain van der Waals pa king. Right, key hydrogen bonds targeting polar threonine and serine residues in TLR4’s TM span. **C)** Non-permeabilized anti-FLAG immunostaining of *de novo* designed LgBit TM proteins show expression and trafficking to the plasma membrane in a predominantly type I orientation upon transfection 293T cells. Scale bar: 10Cμm. **D)** PPI NanoBiT luminescence normalized by LgBit-HiBiT luminescence of LgBiT expression levels. De *novo* TM protein exhibit greater PPI signal upon co-expression with TMJM-SmBit target than with the non-specific control (CD8α-LgBit) (two-tailed *p*-values from student’s t-test), including lead candidates Designs-1 and −6. Representative of n=3 biological replicates. Dots are technical triplicates

For *de novo* design using these template backbone models, we fixed the TLR4 TM domain sequence to one of the TM helices and used Rosetta membrane framework^53,54^ to sample sidechains and guide the selection of synthetic TM adaptor sequences. Amino acid sampling was limited to membrane-compatible amino acids as previously described^9,52,55^. Our 2 guiding design principles were to engage one or more of TLR4’s weakly polar Ser/Thr sidechains in hydrogen bonding (H-bonding) per design and to maximize the number of close backbone-directed knobs-into-holes packing events (<3.0 Å to backbone heavy atom) at the designed TM interface (Figure 2B). We rationalized designs achieving this H-bonding may improve specificity to TLR4’s unique chemistry. The second design goal is driven by recent studies correlating high stabilities in TM domain complexes^46,56^ when apolar van der Waal (vdW) packing achieving this geometric criteria is optimized, reflecting a potential key determinant of how apolar interfaces achieve favorable energetics within the similarly apolar lipid environment. Using iterative rounds of rotamer trials, we sampled hundreds of designs per backbone and ranked those on number of H-bonds, number of tight backbone-directed knobs-into-hole packing events and predicted interface energies. After selection of top designed interface sequences, lipid-facing surface residues were diversified to reflect the natural distribution of membrane-embedded amino acids and remove potential self-interaction sequence patterns. We attempted to use AlphaFold3 structure prediction to assess the likelihood that each synthetic TM peptide will engage TLR4 in the intended binding mode, but predictions were of low quality (low contact density) and dissimilar to the designed TM geometry in all cases (Supplemental Figure 2C) thus not useful to rank designs. Instead, as an orthogonal structural quality metric we used 500 nanosecond all-atom molecular dynamics simulations in POPC bilayers to assess the interface stability of the top TM designs in complex with TLR4’s TM span. We found 9 unique designs passed stability criteria of retaining <2 Å mean backbone RMSD from the initial model (Supplemental Figure 2B). This computational pipeline is accessible to anyone with a single-GPU workstation—or even a laptop, if molecular dynamics (MD) simulations are skipped. A single round of design rapidly generates candidate transmembrane interaction sequences in high quality models passing our *in silico* metrics, stable in MD, by optimizing features we propose as design principles for recognizing predominantly apolar TM spans and suited for test the intended function to bind and antagonize TLR4 signaling.

To assess whether these top 9 *de novo* TM polypeptides (Supplementary Table 1) bind TLR4’s TM domain in mammalian cell membranes, we used the adapted NanoBiT assay to evaluate protein-protein interactions upon co-expressing LgBit-fused “Binders” with the TLR4-TMJM-SmBit “Target”, again comparing their relative luminescence to that of baseline complementation from co-expressing non-interaction negative control TM proteins (Supplemental Figure 2D). Fluorescence microscopy staining of cell surface FLAG-tag without permeabilization confirmed robust expression and plasma membrane localization of the *de novo* TM proteins consistent with the expected Type I insertion topology when co-transfected with TLR4-TMJM-SmBit, matching parallel-oriented helices as designed (Figure 2C). In the NanoBiT assay, 5 of the 9 designed TM proteins exhibit much higher normalized luminescence signal than the non-interacting CD8α control, indicating potential TLR4 binding complexes (Figure 2D, Supplemental Figure 3). Design-1 and Design-6 demonstrated the highest interaction propensity with TLR4-TMJM-SmBit, significantly greater than that of TLR4 TMJM’s self-interaction by 1.6±0.1-fold and 2.0 ± 0.2-fold normalized luminescence, respectively (student’s t-test; two tailed p-value <0.05 and <0.01 respectively). Thus, amongst a high overall apparent success rate of TM domain PPI design, these two designed proteins were identified as lead candidates for anti-TLR4 binding and further evaluated for potential functional regulation.

Reviewing the interface features in these 2 putative successful TLR4-binding designs reveals they both span a common binding region of TLR4 and share similar optimized features prioritized in design. Design-1’s backbone binding pose was a TMHop-derived prediction; its design model achieves 12 tight backbone-engaging knobs-into-holes packing interactions with TLR4. Design-1’s 1 mid-spanning polar residue, Thr13, was intended to target TLR4’s S640 via hydrogen bond, but in MD simulations Thr13’s sidechain instead favors an intra-helical H-bonding conformation that packs its methyl group against TLR4 (Figure 3A). Consequently, Design-1 primarily relies on apolar vdW interactions for its recognition and complex stability with TLR4. Design-6’s backbone binding pose was derived from an AlphaFold2-predicted model and is expected to achieve interface stabilization via a combination of H-bonding and apolar packing: 8 backbone-engaging knobs-into-holes events and a stable inter-chain H-bond from Design-6’s Thr13 to TLR4-TM’s Ser640 (Figure 3B). Design-1 and Design-6 adopt distinct parallel inter-helical crossing angles and target divergent sections of TLR4’s C-terminal TM domain. However, they both engage the similar N-terminal helix face of TLR4 bearing T632, V636, S640, V643, V647 – residues whose chemical shifts were most sensitive to oligomerization in solution NMR studies and is thus the most likely lowest energy TM dimerization interface^35^.

**Figure 3.**
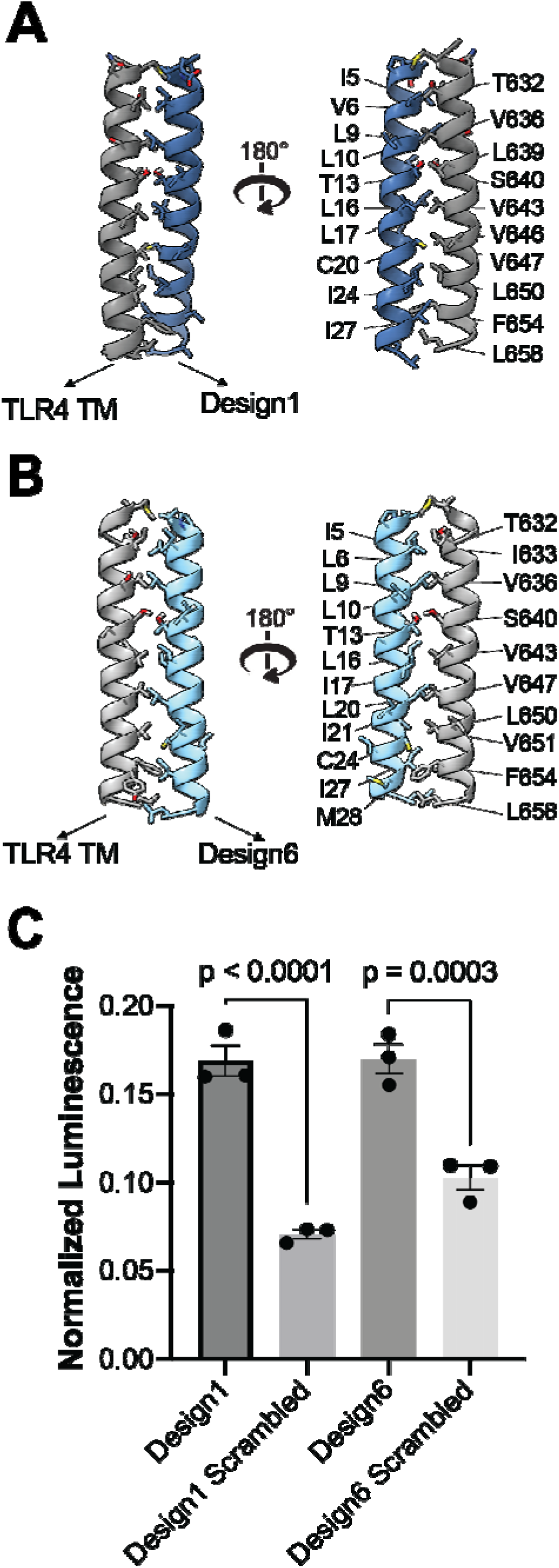
Interactions between TLR4 TMJM and Designed TM proteins are sequence-specific. A) Structural model of anti-TLR4 Design-1 (dark blue) bound to the TLR4 TM domain (grey), mediated primarily by packing of apolar interfacial side chains. B) Structural model of anti-TLR4 Design-6 (light blue) bound to the TLR4 TM domain (grey), highlighting key T13-S640 inter-helical hydrogen bond alongside optimized apolar knob-into-hole interface sidechain packing. C) Scrambled sequence variants of Designs-1 and −6 exhibited significantly reduced expression-normalized NanoBiT luminescence relative to their respective original designs upon co-expression with TLR4-TMJM-SmBit, reflecting sequence-specificity. Representative of n=3 biological replicates. Dots are technical triplicates. Error bars, SEM. P-values from two-tailed student’s t-test.

To test sequence specificity of these lead anti-TLR4 TM peptides, we first generated modified constructs scrambling the TM span primary sequences and tested their interaction with the TLR4-TMJM fragment (Supplemental Figure 4A). Scrambled Designs-1 and −6 exhibited significantly lower luminescence upon co-expression with TLR4-TMJM at levels similar to non-interacting control TM proteins (e.g. CD8α), much lower than non-scrambled designed proteins (2.5**±**0.5 fold-change, p = 0.002; 1.8**±**0.4 fold-change, p=0.045; both n=3; Figure 3C, Supplemental Figure 4B). This apparent disruption of the protein-protein interaction supports that the specific sequence features of Designs-1 and −6, engineered by computational design, are required to engage TLR4—rather than inherent amino acid compositional properties such as TM length or hydrophobicity, which may also influence membrane microdomain partitioning or bilayer crowding effects.

Next, we assessed the specificity of Design-1 and −6 for TLR4 versus a small panel of alternate SmBit-tagged membrane proteins (tetraspanin CD81, GPCRs mu-opioid receptor and β2-Adrenergic receptor, and single-pass surface receptors EGFR and EPOR). The two designs have the strongest NanoBit PPI signal with TLR4’s TMJM fragment and have essentially no interaction propensity above baseline split enzyme complementation with 4 of 5 alternative membrane proteins (i.e. less than or equal to that of non-interacting control protein CD8α with each SmBit-tagged “Target”; Supplemental Figure 4C). Design-6 exhibits a strong off-target PPI with muOR, while Design-1 interacts with muOR but much less so (∼3-fold) than it does with TLR4-TM. While the mechanism of promiscuous interaction with muOR is unclear, this finding represents a limitation in our computational targeting and highlights the potentially broader challenge of engineering apolar TM peptides for specificity spanning membrane proteins of the cell. Future studies underlying the nature of this interaction and refinement of the chemical approach (i.e. negative design) may be needed to use the probes in more complex phenotypic interrogation or as potential therapeutics.

### Biological Activity of De Novo Designed TM Polypeptides

We next investigated whether the apparent association of synthetic TM polypeptides with TLR4’s TM span is sufficient to disrupt full-length receptor signaling in mammalian cells. To do so, we transiently transfected LgBit-fusion Design-1 and −6 alongside TLR4-eGFP into the NF-κB-SEAP reporter assay and challenged with LPS stimulation. Upon saturating LPS stimulation of TLR4 signaling, we find co-expression of Design-6 and Design-1 significantly reduces the maximal response of TLR4, (i.e. signaling efficacy), corresponding to 46.7_±_3.0% and 35.1_±_7.4% inhibition, respectively (n=3; both p <0.05; Figure 4A,B). The scrambled variant of Design-1 also suppresses TLR4 signaling efficacy (26.6_±_10.9%, n_=_3, p < 0.05), whereas the scrambled Design-6 demonstrates negligible antagonism of LPS-induced receptor signaling (3.54_±_9.1%, n_=_3). These LgBit-tagged TM-targeting proteins are all well expressed at comparable levels, reflecting relative activities are determined by their specific sequences (Figure 4D, Supplemental Figure 1C, Fig 2C). Given Design-1’s scrambled variant is sufficient to inhibit signaling despite not associating with TLR4, both its mechanism and that of Design-1’s inhibition are questionable – possibly via indirect consequences and not via means of binding. Co-expression of the designed TM proteins have minimal effect on TLR4 basal receptor signaling and Design-6 expression has minimal impact of TLR4 protein expression level, ruling out the alternative sources of reduced signaling resulting from functional TLR4 levels upon co-expression (Figure 4A,C, S5B). Design-6’s inhibitory effect is dose-dependent, correlates with apparent protein levels, and saturates at higher expression conditions (Supplemental Figure 5). Furthermore, Design-6 reduces the maximal signaling response without shifting the EC__, indicating it does not interfere with ligand binding or receptor sensitivity. Thus, Design-6’s appears to encode specific cross-membrane signaling antagonism via its putative TLR4 complex inhibition, which we hypothesize results from restricting the receptor’s access to its signaling-competent TM conformation within the membrane (Figure 4A).

**Figure 4.**
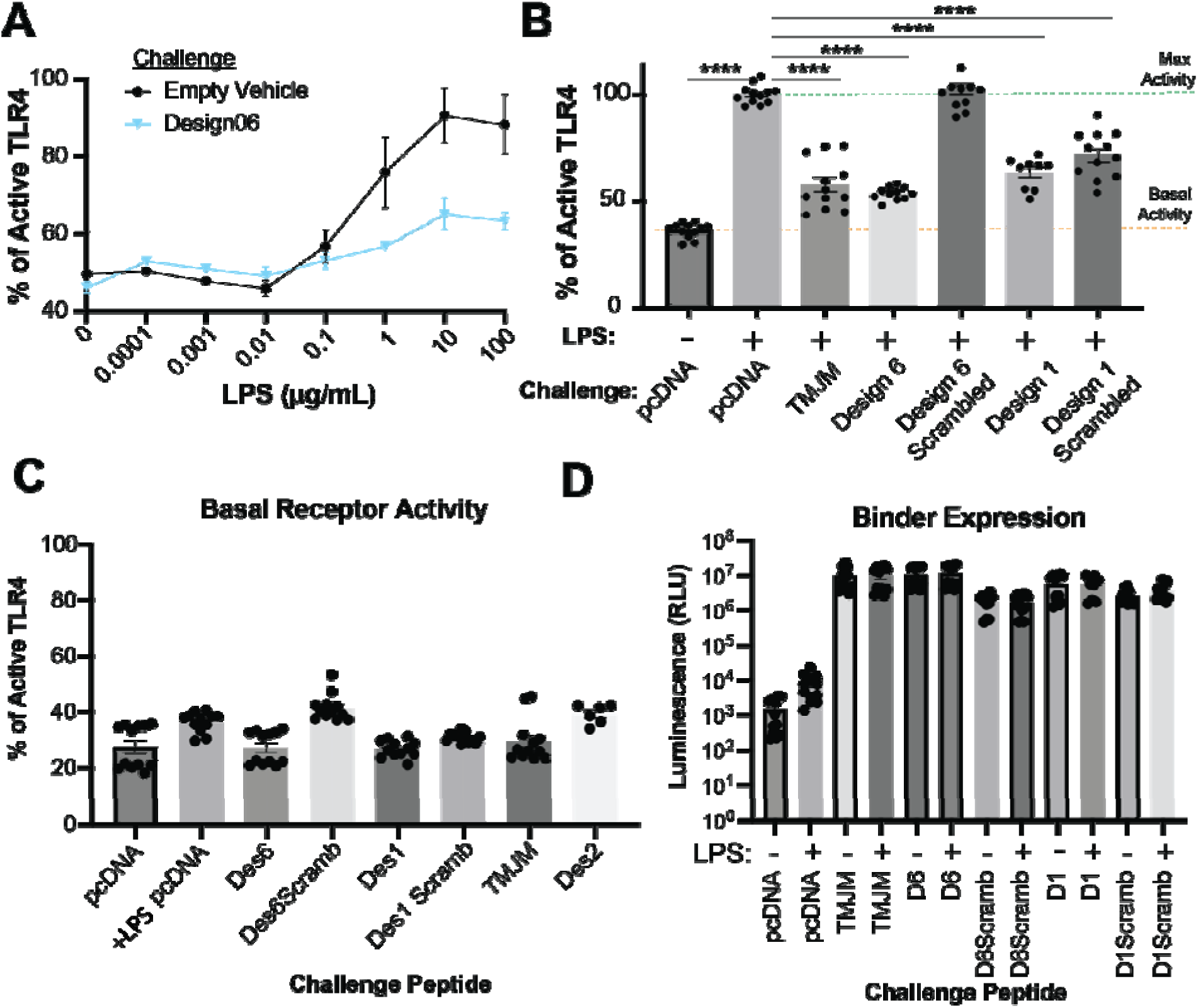
Designed TM proteins inhibit cellular TLR4-mediated NF-_κ_B signaling activation. A) LPS dose-response curves of TLR4-eGFP co-expressed with Design-6 (cyan) show reduced total signaling response compared to TLR4-eGFP co-transfected with empty vector (black), indicating a non-competitive inhibitory mechanism. Dots are technical triplicates for n=3 biological replicates. Error bars, SEM. Aster sks denote two-tailed p-value of significance by student’s t-test (**** p≤0.0001). B) NF-κB reporter signal in cells without (untreated, basal signaling) or with (10 µg/mL, max activity) LPS stimulation after co-transfection with TLR4-eGPF and plasmids encoding empty vector, TLR4 fragment TMJM, lead TM protein designs, or their scrambled sequence controls. Relative to maximum activity of LPS-stimulation of TLR4 signaling alone (green dot line), Design-6 inhibits NF-κB activation similarly to TLR4-TMJM while Design-6-scrambled has no effect. Design-1 and its scrambled variant showed moderate inhibition. Dots represent technical replicates from n=3 biologically independent experiments. Error bars, SEM. Asterisks, p-value <0.0001 in t-test using n=12. C) Basal NF-κB activity in experimental conditions for cells expressing TLR4 alone (pcDNA) or co-transfected with TM proteins in the absence of LPS stimulation. Dots represent technical replicates from n=3 biologically independent experiments. Error bars, SEM. D) Luminescence upon addition of HiBit peptide measuring LgBit-fused protein expression levels show protein copy number is comparable across controls and lead TM protein designs co-transfected in 293T cells with or without LPS (n = 3 biological replicates).

### Design 6 and TLR4 TMJM form a stable biochemical complex

To test whether the observed antagonism of TLR4 signaling is a result of a stable and direct biochemical interaction with Design-6, we first characterized the complex extracted from cell membranes. Co-immunoprecipitation (co-IP) was used to test formation and stability of the membrane-associated protein interaction with TLR4 from the mammalian cell environment and upon detergent extraction. The membrane fraction of HEK293T cells expressing HA-tagged TMJM target were pulled down by co-expressed FLAG-tag bait construct Design-6, TMJM, or eVgL solubilized by 1% n-dodecyl-β-D-maltoside (DDM) and 0.1% cholesterol hemisuccinate (CHS). Immunoblotting for HA revealed significant enrichment of the TLR4-TMJM protein upon co-IP by Design-6 but much less so by a homotypic TLR4-TMJM fragment or non-interacting control protein eVgL (Supplemental Figure 6A). Thus, Design-6 forms a stable, detergent-extractable complex with TLR4’s TM region in human cells, corroborating that direct interactions mediate their cellular proximity measured NanoBit split enzyme complementation.

Next, to further investigate the nature and stability of Design-6’s interaction with TLR4’s TM segment, we measured the ^15^N-^1^H HSQC spectra of a U-^15^N TLR4-TMJM protein fragment in ^2^H-DPC micelles with or without Design-6 as a minimal TM peptide by solution NMR (Figure 5, Table S1). To distinguish spectral changes due to Design-6 binding versus potential detergent-dependent oligomerization of TLR4-TMJM, we first measured spectra of TLR4-TMJM alone upon varying dodecylphoshocholine (DPC) concentration, titrating the detergent:protein molar ratio at 600x, 180x, and 60x (Fig S6B,C,E). Between 600x and 180x, TLR4-TMJM’s ^1^H-^15^N HSQC spectra exhibits extensive chemical shift perturbations and broadening in apparent fast exchange, likely reflecting homo-dimerization events as seen in previous NMR studies^28^. At 60x (∼1 protein per micelle), most TLR4-TMJM peaks further broaden, and many undergo additional perturbation but in different directions from those seen between 600x and 180x spectra. This behavior indicates that TLR4-TMJM likely reaches a mixture of oligomers beyond physiological homodimers, also previously observed previously at these near-crowding conditions^28^. When TLR4-TMJM is reconstituted with Design-6 at increasing molar equivalents, its resonances are broadly perturbed relative to spectra of TLR4-TMJM’s alone both at 180x (likely homo-dimeric) or 600x (likely monomeric) conditions (Fig 5, S7). Given the unique induced perturbations and distinct chemical environment compared to TLR-TMJM monomer-oligomer equilibria, these spectra indicate Design-6 forms a stable binding complex in these relative harsh conditions: 45*°* C in zwitterionic detergent. Design-6 binding appears sharpen resonances, not inducing further broadening relative to 180x DPC of TLR4-TMJM (likely homodimer state). These apparent relative dynamics combined with the distinct peak shift directionalities versus the TLR-TMJM monomer upon peptide titration suggest Design-6 is likely binding the TM monomer, rather than associating into larger TM assemblies with the already more dynamic TLR4 oligomeric species (Fig S7A,F). Likewise, observing the Design-6/TMJM complex is favored under conditions where TLR4-TMJM is otherwise homodimeric suggests Design-6 interacts competitively with TLR4-TMJM homo-dimers (Figure S7). Competitive Design-6 binding suggests its interactions may lie on the same TM helix surface overlapping with TLR4’s homo-dimerization interface, being mutually exclusive. Collectively we find a direct and stable interaction complex of Design-6 with TLR4 can be isolated *in vitro,* indicating our computational strategy successfully engineers membrane-embedded interactions. The designed membrane adaptor complex that readily assembles *in vitro* is most likely responsible for its antagonism of TLR4 NF-κB signaling in mammalian cells as the mechanism of synthetic immune regulation.

**Figure 5.**
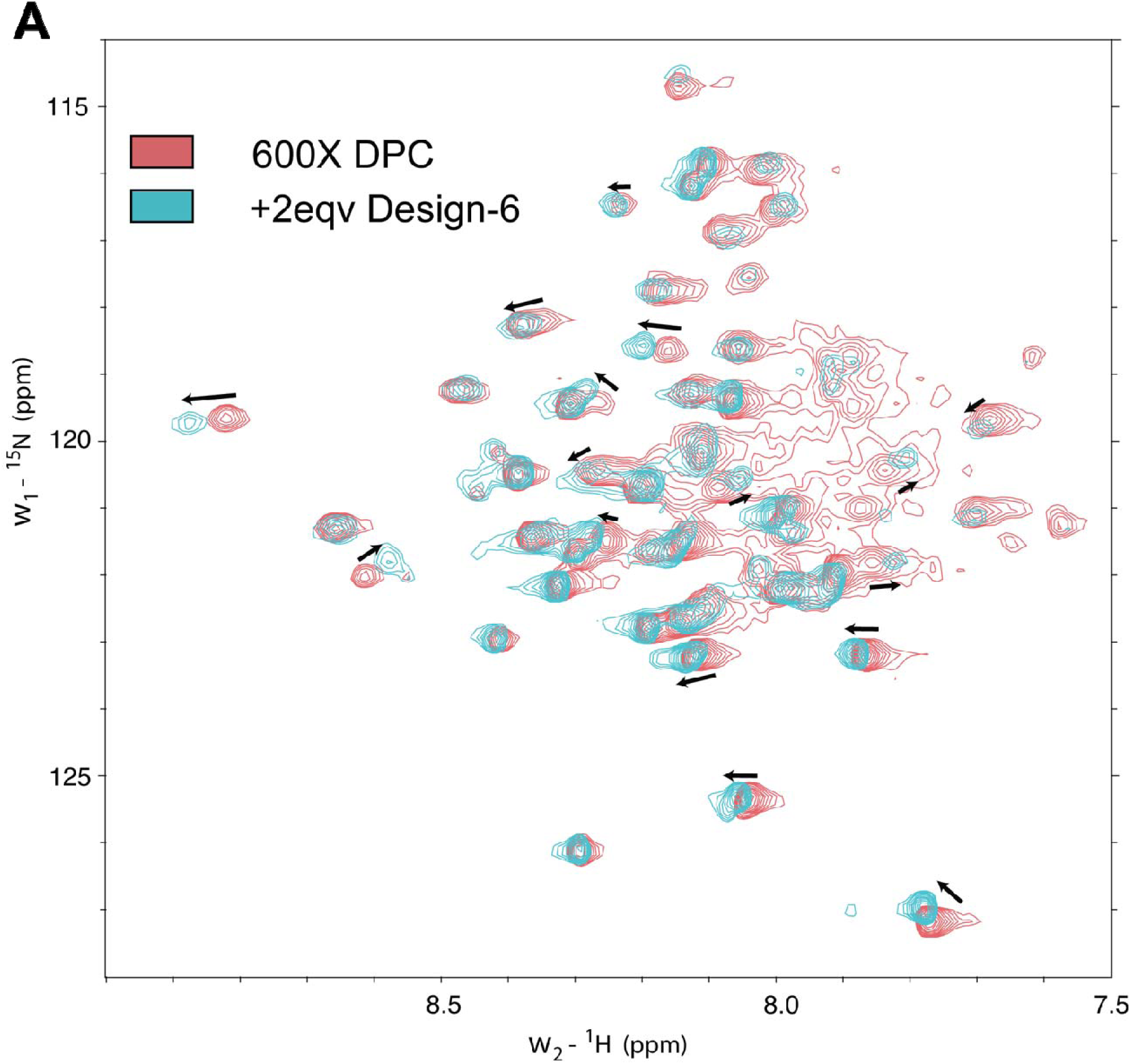
Solution NMR of TLR4-TMJM complex with Design-6 in DPC micelle. The ^1^H-^15^N HSQC spectra of 0.2CmM ^15^N TLR4-TMJM at 40C°C, pH 6.0, and 800CMHz, either with TMJM alone in ^2^H-DPC at 120 mM representing the monomeric state (red, 600:1 detergent:TMJM molar ratio, red) or with 0.4 mM Design-6 in 36 mM ^2^H-DPC (blue, 180:1 detergent:TMJM; 60x detergent to total protein molar ratio). Arrows denote chemical shift perturbations. Eqv, equivalents.

## DISCUSSION

Our successful demonstration of molecular design targeting this apolar TM span of an immune receptor without prior structural templates extends the forefront of technological capacity for engineering sequence recognition and building stabilizing interactions within lipid bilayers on-demand. We devised new working principles for selecting stabilizing heterotypic apolar interfaces which led to a high success rate (>10%) relative to similar cutting-edge accomplishments in protein design, supporting our hypothesis that optimizing backbone-directed sidechain packing is a viable route to effectively rank design molecular models and encode stability in TM assemblies^46,56^. Bolstered by this structural quality metric, we establish this membrane protein design framework as suitable to expand molecular targeting to engage residue compositions typical of most single-pass TM domain in the proteome – not just the fraction of TM spans with small residue motifs. These design principles and filtering metrics can be integrated with AI/ML approaches to model new binding sites (RFdiffusion^57^) and sample potential sequences (MPNN^58^) for greater customization, not limited to symmetric homodimer prediction templates and possibly adaptable to multi-spanning proteins. The computational methods proven in practice here catalyze generation of new functional chemical matter and thus enable probing of novel transmembrane domain interaction mechanisms of other Toll-like receptors and diverse membrane protein families or immune complexes.

TLR4 antagonism observed with natural and designed TM domain polypeptides reveals new insights into cross-membrane signaling and gives rise to further opportunities to interrogate the mechanism for therapeutic benefit. Given TLR4 constructs lacking the ectodomain are constitutively active, it follows that when given geometric freedom the conformations preferred by the TM-JM and cytoplasmic TIR domains are cooperative and productive for signaling^30^. In the full-length receptor, the ectodomains likely oppose the driving force towards activation from the underlying modest strength TM dimer, effecting regulatory autoinhibition via general steric occlusion of the large domain or specific structural restraints from signaling-incompetent monomeric and dimeric conformations. Our findings that disrupting TLR4’s lipid-embedded dimerization impairs signaling reveals that achieving a particular TM domain geometry is a key determinant for cross-membrane coupling, rather than the TM region being a passive anchor shifted freely upon ligand binding and dimeric ectodomain rearrangements. We propose a model where ligand chemistry tunes the energetic balance and geometric coupling of these domains, aligned with our data and recent cryo-EM structures^1^. Precise conformational changes relieve autoinhibition and appear to determine receptor signaling efficacy by shaping how the ectodomain geometry connects to the transmembrane region. Thus, the interplay between resulting TM geometries and effectiveness in inducing downstream signaling is an emerging feature to leverage in the design of new therapeutic moieties. How TLR4 ligands act to bias signaling towards interferon-stimulated genes versus the NF-κB pathway is an outstanding question of paramount biomedical importance. Given our observations, the role of the TM domain becomes of increasing interest in TLR4 bias; new TM-directed probes could serve to dissect this behavior. Furthermore, TLR signaling also includes alternative mechanisms beyond direct ligand-induced conformational changes that impact signaling: receptor clustering, receptor internalization, signaling in intracellular compartments (e.g. endosomes), and regulatory post-translational modifications, namely palmitoylation^59^. Given the involvement of lipid bilayers in each of these behaviors, it is possible TM domain features play an active role and TM-directed modalities chemically modulating these activities could help clarify more thorough models of TLR4 signaling.

The molecules and strategy for *de novo* design demonstrated here add to the growing toolkit and precedent of intercepting immune signaling within the lipid bilayer for mechanistic insights and therapeutic potential^6,49^. TLR4 overactivity is coincident in many inflammatory disorders and thought to be causal in colitis and sepsis, making this receptor a prime but underutilized target in autoimmune intervention – lacking effective drugs (i.e. no FDA-approved antagonists). Given difficulty for targeting at large apolar lipid-binding surfaces of ectodomain orthosteric sites and complexity of modifying lipid endotoxin scaffolds, our ability to generate active antagonist TM polypeptides provides an interesting novel route to regulate and probe TLR4 modulation for potential benefit in models of disease. While established here in a canonical inflammatory signaling HEK293T cell model, future work should address activities of these peptides for biasing signaling pathways and functioning in more physiologically relevant immune cell types. Another boundary to therapeutic application of computationally designed TM sequences is delivery, given their highly hydrophobic nature. Previous work leveraged limited solubility of minimal TM peptides to penetrate cells and tissues by spontaneous membrane insertion^8,9,48,60,61^. There is now growing opportunity and precedent for delivery of these sequences either genetically (e.g. mRNA, using the cells’ own translational machinery) or as peptide derivatives; both could benefit from lipid nanoparticle delivery systems as seen previously^12^. Chemical targeting of membrane proteins directly within the lipid bilayer is still in its infancy. The advancements in design of new lipid-embedded molecules with impactful biological consequences reaching a broader protein scope paired with emerging strategies for efficacious delivery can enable evaluation of novel TM-direct strategies for therapeutic membrane protein modulation.

