## Supplemental Figures for "Programmed inhibition of an innate immune receptor via *de novo* designed transmembrane proteins"

| Design 1 | EAPLIVWLLLLITIFLLGICAFIIYLIKKLA |
| --- | --- |
| Design 2 | EAPLIVWLLLLITIFLLGIMAFIIYLMKKLA |
| Design 3 | EEPLIVLALIGIISLLIFIMFGIIWLMKK |
| Design 4 | EAPELWLILGLILAMILILTLLIATMKKLLG |
| Design 5 | EAPLLCWLNILITIFLIATLLLIMLKR |
| Design 6 | EAPLILWLLLGITAFLILLLIGICLLIMKKL |
| Design 7 | EAPLINWLIILITIFIIGTLILIIKK |
| Design 8 | EEPEVTLWITALCIMAILLITGVLFYLMKKD |
| Design 9 | EAPLIMWLAILITIFIIGLLIFIMLALLKKL |
| eVgL | DPGSLWAIVFLLFLIVLLLLAIVFLLRR |
| CD8a | FVLTLSDFRRENEGYYFCSALSNSIMYFSHFVPVFLPAKPTTT  PAPRPPTPAPTIASQPLSLRPEACRPAAGGAVHTRGLDFACDIY  IWAPLAGTCGVLLLSLVITLYCNHRNRRRVCKCPRPV |
| TLR4-TM | MNKTIIGVSVLSVLVVSVVAVLVYKF |
| TLR4-TMJM | MNKTIIGVSVLSVLVVSVVAVLVYKFYFHLMLLAGCIKYGRGE |
| Design-6 (NMR) | EEEEAPLILWLLLGITAFLILLLIGICLLIMKKLRRK |
| TMJM (NMR) | MHHHHHHGSDSERERPGLVPRGSADHDAP  ^628^QMNKTIIGVSVLSVLVVSVVAVLVYKFYFHLMLLAGCIKYGRGEN^672^HEWRE |

**Supplementary Table 1. Transmembrane domain sequences of natural TLR4 fragments and *de novo* designs**

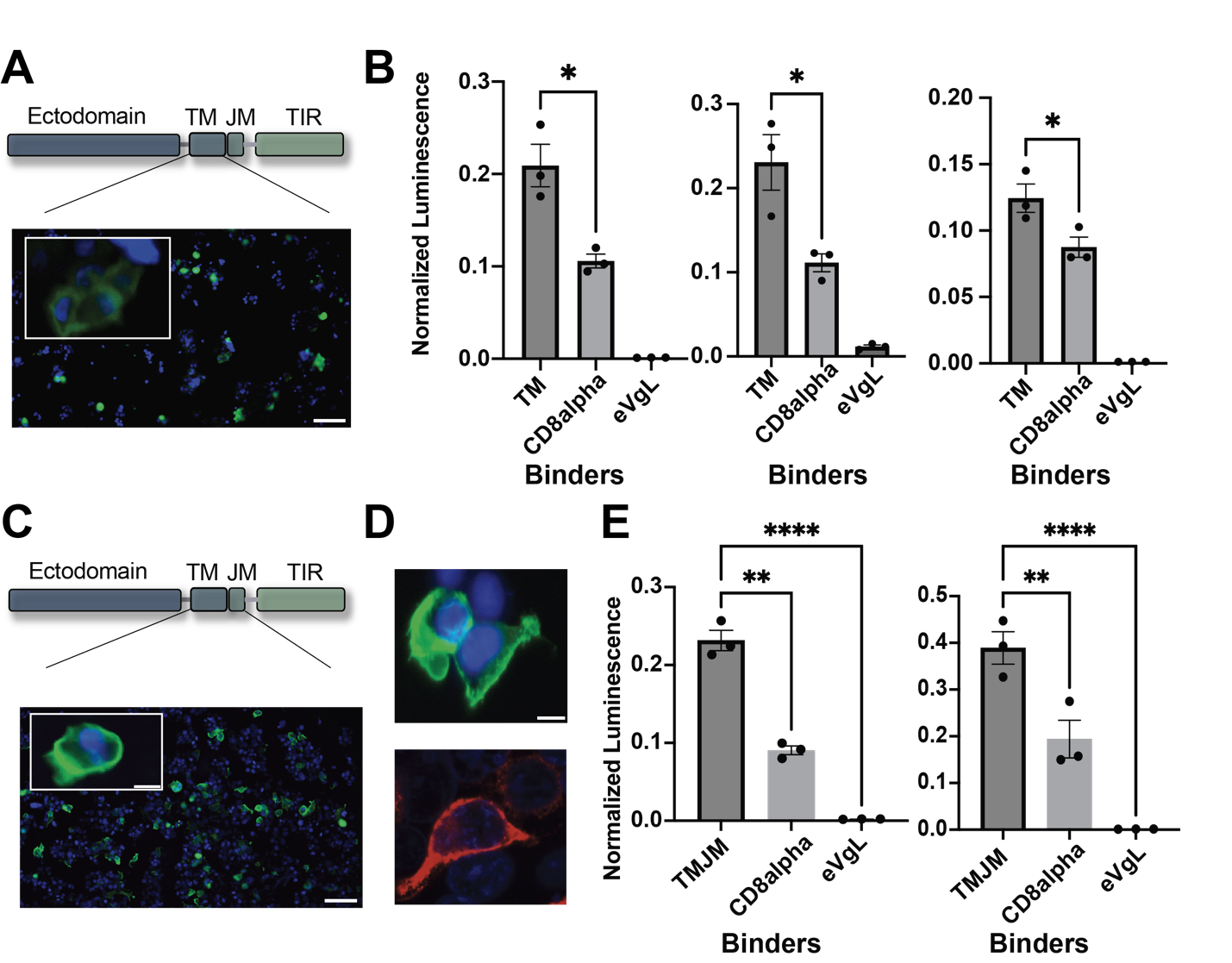

**Supplemental Figure 1. Membrane spanning domains of TLR4 self-associate in mammalian cells**

A) Immunostaining of TLR4-TM-lgBit “binder” construct expressed in 293T cells and labeled shows moderate expression and trafficking to the plasma membrane. Scale bar (widefield): 100 μm. Scale bar (inset): 10 μm.
B) Individual biological replicates of NanoBiT luminescence (SmBiT–LgBiT interaction) from Figure 1B normalized by subsequent LgBit-HiBiT luminescence (LgBiT expression), showing that TLR4-TM-SmBit co-expression leads to stronger relative split enzyme complementation with TLR4-TM-LgBit than with than with non-specific negative control proteins, CD8α-LgBit and eVgL-LgBit. Dots represent technical triplicates. Error bars, SEM. Asterisks denote two-tailed p-value of significance by student’s t-test (* p≤0.05).
C) Immunostaining of the TLR4-TMJM-LgBit “binder” protein with anti-Flag–Alexa488, shows slightly higher expression and more efficient plasma membrane localization compared to TLR4-TM-LgBit.
D) Immunostaining of the CD8α (top, anti-Flag-AF488) and eVgL (bottom, anti-FLAG-AF647) confirms comparable expression and trafficking.
E) Additional biological replicates of NanoBiT assay as in Figure 1B (n=3 total), demonstrating that TLR4-TMJM-SmBit paired with TLR4-TMJM-LgBit interaction results in an approximate 2-fold increase in normalized luminescence compared to CD8α-LgBit co-expression and >100-fold increase relative to eVgL-LgBit co-expression. Dots represent technical triplicates. Error bars, SEM. Asterisks denote two-tailed p-value of significance by student’s t-test (** p≤0.01; **** p≤0.0001).

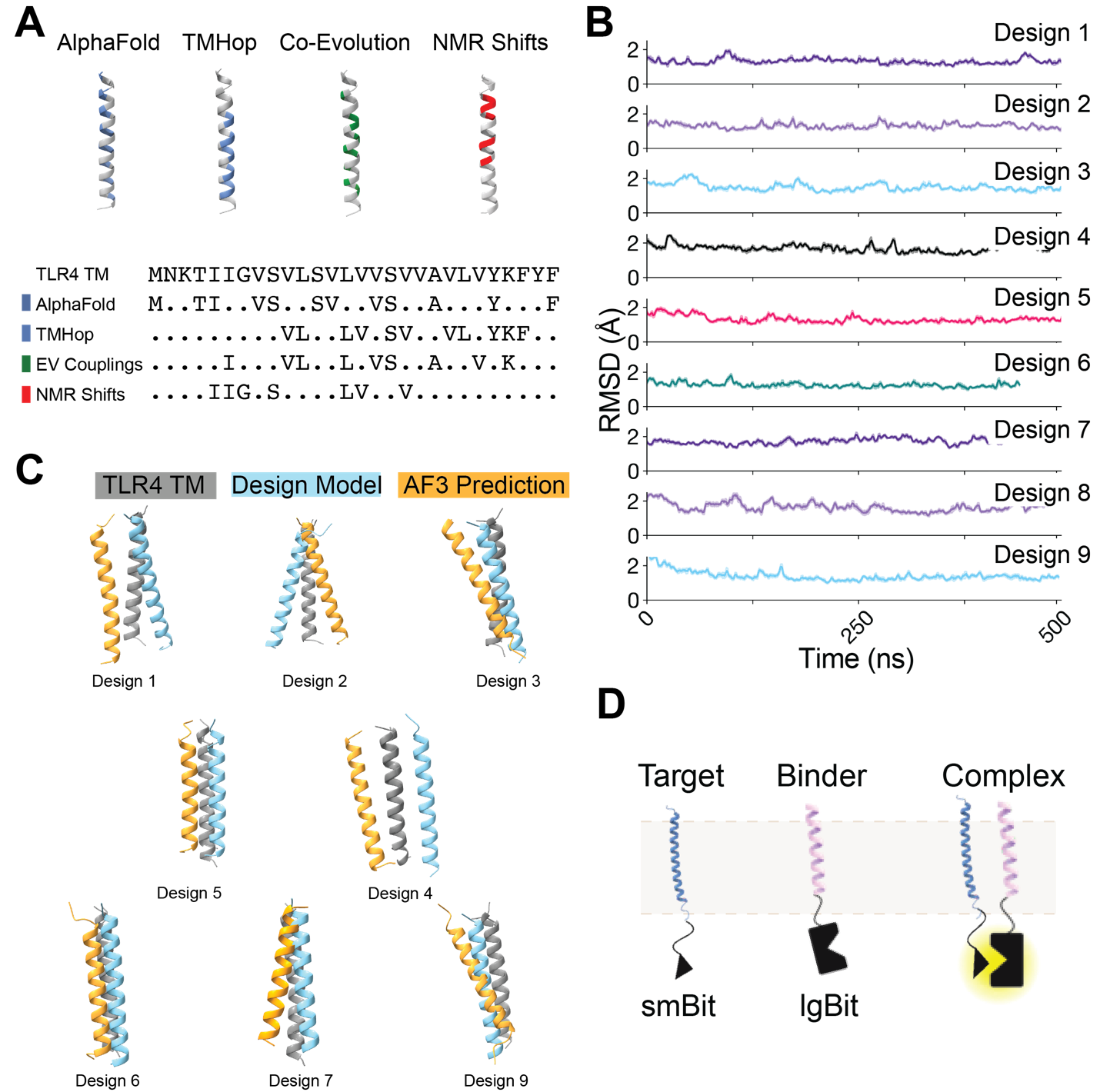

**Supplemental Figure 2. In silico and in cell validation strategies for TLR4 targeting designs**

A) Top, predicted dimer interface residues colored on cartoon TM helix by prediction source (blue: AlphaFold2, TMHop; green: EVCouplings; red: NMR shifts). Bottom, TLR4 TM domain sequence and residues predicted at dimer interface by each approach, revealing no convergence to a clear TM domain dimer geometry.
B) All-atom molecular dynamics simulations in model POPC membranes of top 9 *de novo* designed TM proteins in complex with TLR4 TM domain exhibit stability, defined by mean backbone RMSD values <2 Å during 500 nanoseconds.
C) AlphaFold3 predictions of complex between TLR4-TM (gray) and each de novo design TM domain (orange) have major discrepancies in TM interface helix geometry compared to the design TM domain binding pose in design models (blue).
D) Schematic of membrane protein-protein interaction assay with split NanoLuc enzyme complementation: TLR4-TMJM fused to the smBit and *de novo* designed TM peptides fused to the lgBit tested in Figure 2.

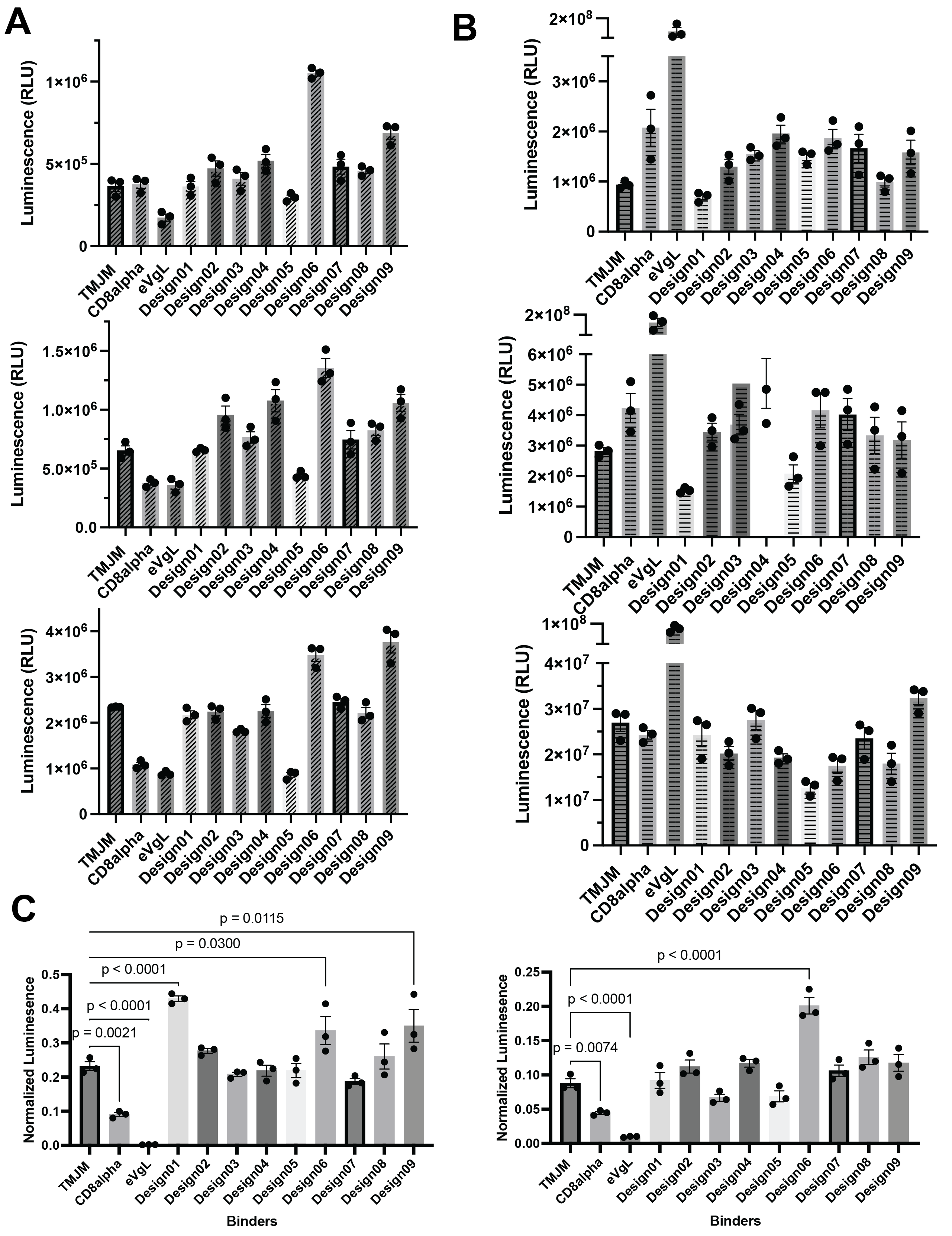

**Supplemental Figure 3. Quantification of transmembrane protein interactions between *de novo* designs and TLR4 TMJM in cells using NanoBit assay.**

A) Unnormalized NanoBiT luminescence resulting from PPI-driven complementation of smBit and lgBit fragments for 3 biological replicates, reflecting interaction propensity between *de novo* designed binders and the TLR4 TMJM target in HEK293T cells upon co-expression. Dots represent technical triplicates of each biological replicate. Error bars, SEM.
B) Secondary LgBit-HiBiT luminescence upon delivery of HiBit peptide to cells quantifying protein relative abundance of each LgBit-binder construct for 3 biological replicates alongside NanoBit PPI measurements in panel A (rows are paired experiments), used to normalize and yield relative luminescence of interaction (n=3 biological replicates with dots representing technical triplicates).
C) Normalized luminescence values of two additional biological replicates (n=3 including Figure 2D) for the 9 *de novo* designed TM LgBit “binders” co-expressed with the TMJM-SmBit “target”, indicating relative interaction capacities. Error bars, SEM. Asterisks denote two-tailed p-value of significance by student’s t-test.

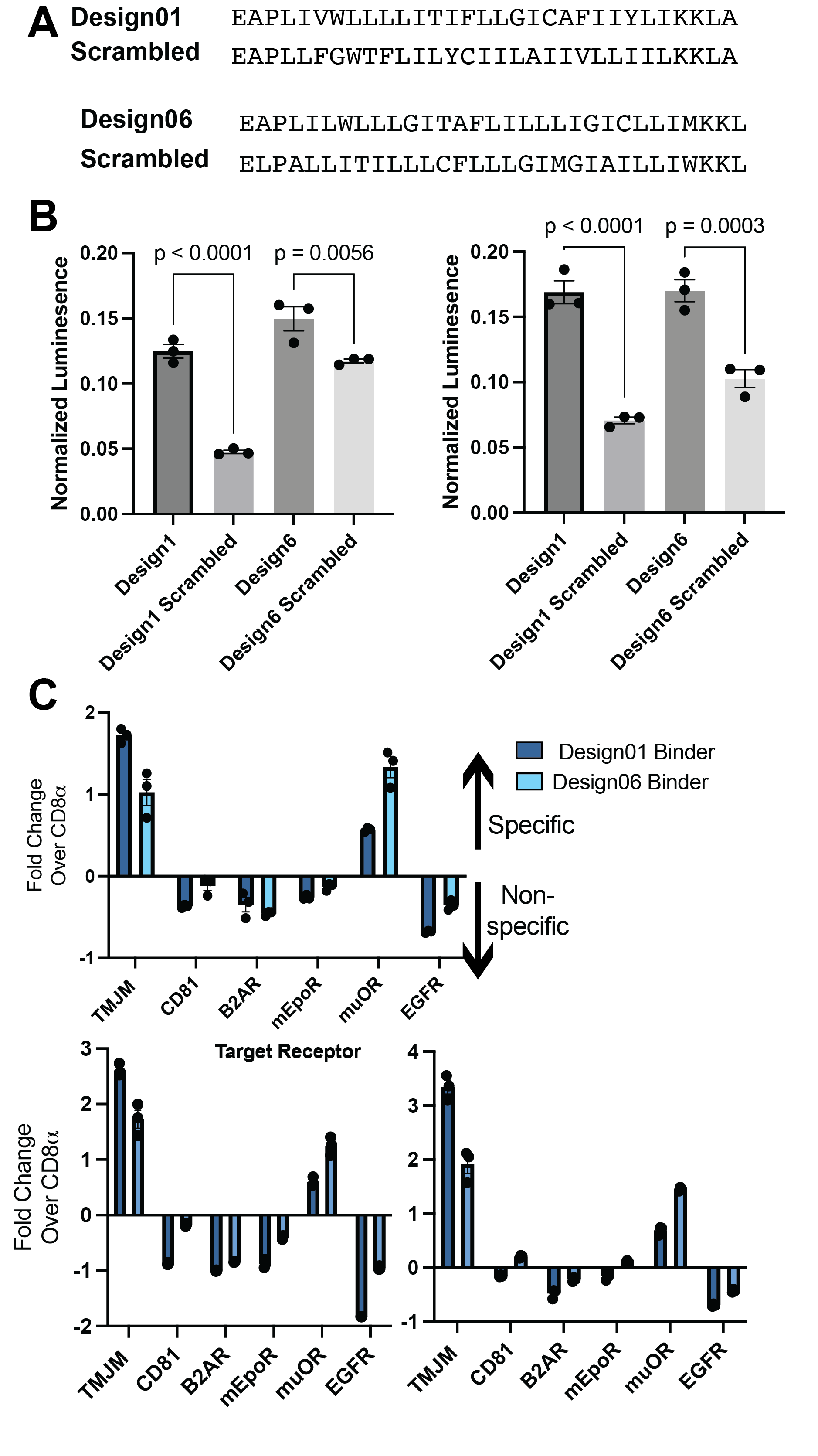

**Supplemental Figure 4. Sequence specificity and target selectivity of lead *de novo* TM binders**

**A)** Designed original and scrambled TM domain primary sequences share < 80% sequence identity.

**B)** Additional biological replicates (n = 3 total with Figure 3B) of expression-normalized NanoBiT PPI assay co-expressing TLR4-TMJM-SmBit with lead Designs-1 and -6 or their scrambled sequence variants, plotted as in Figure 3B.

**C)** NanoBiT interaction of lead LgBit designed TM proteins co-expressed with a panel of SmBit-fused natural single- and multi-pass membrane proteins (n = 3 independent biological replicates, dots are technical triplicate, error bars, SEM). Data are expressed as fold-change relative to the normalized PPI signal of each SmBit membrane protein co-expressed with the non-specific control CD8α, where values >0 indicate specific interactions and values <01 indicate non-specific binding.

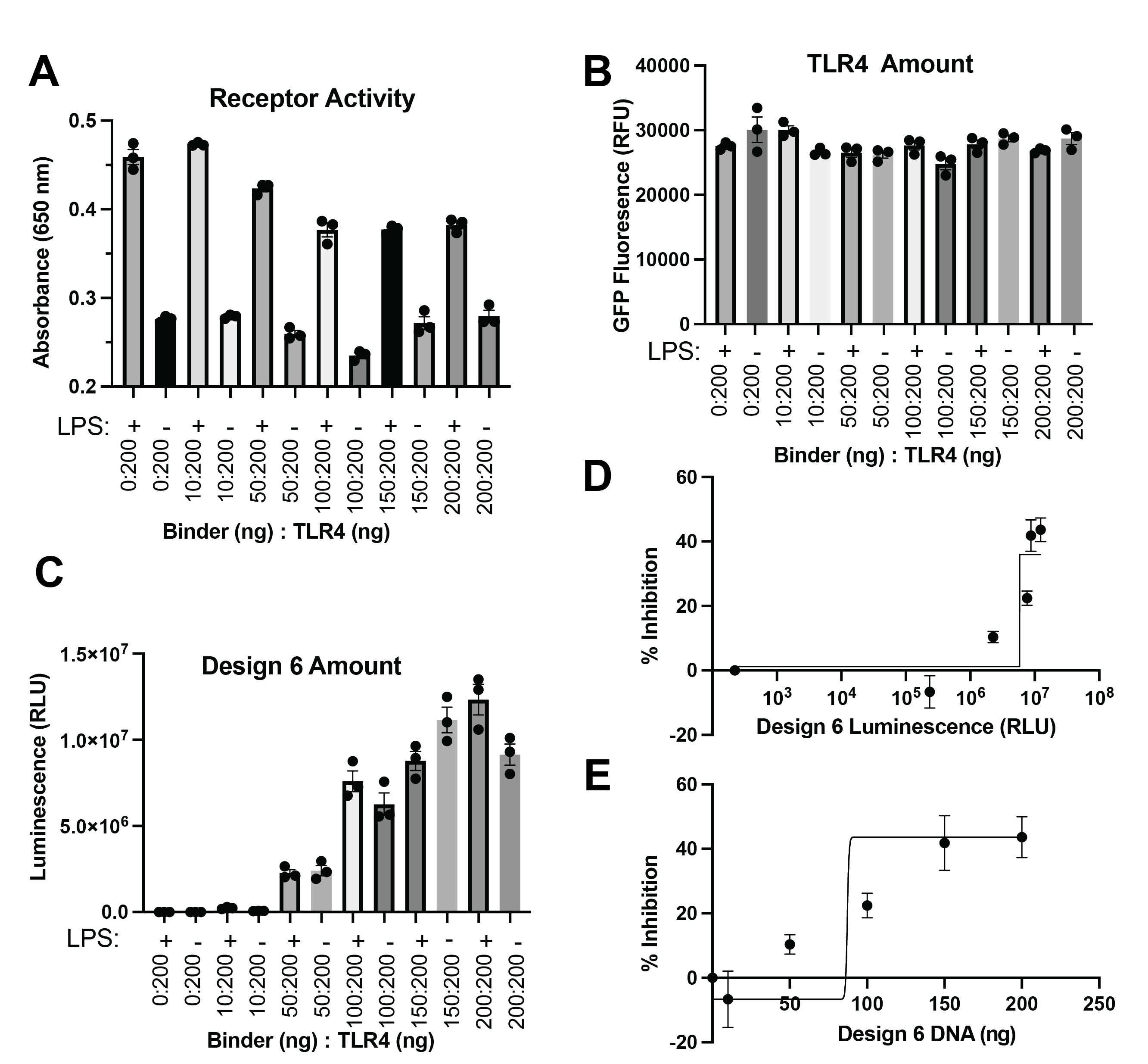

**Supplemental Figure 5. Relative expression levels and Design-6 Dose Dependent TLR4 inhibition**

A) Raw absorbance of TLR4 signaling activation in HEK-Blue NF-κB reporter cells with TLR4 decreases progressively with increasing amounts of Design 6 plasmid co-transfected upon stimulation with 10 µg/mL LPS. Dots are technical triplicates from n=1 experiment.
B) TLR4-GFP fluorescence intensity when co-expressed with Design-6 with (10 µg/mL) or without LPS treatment is not significantly different across varying co-transfection conditions (ANOVA, p > 0.5), indicating that receptor degradation is not a significant contributor to the observed inhibition by lead designs.
C) Luminescence upon addition of the HiBit peptide confirms a corresponding increase in Design-6-LgBit expression proportional to the DNA ratio transfected with pcDNA in 293T cells co-expressing TLR4, with (10 µg/mL) or without LPS.
D) Percent inhibition of NF-κB reporter normalized to condition’s basal activity as the amount of Design-6 increases, plotted as a function of LgBit-HiBit expression level luminescence (RLU) for Design 6.
E) Percent inhibition of NF-κB reporter normalized to condition’s basal activity as the amount of Design-6 expressing plasmid is delivered in co-transfection.

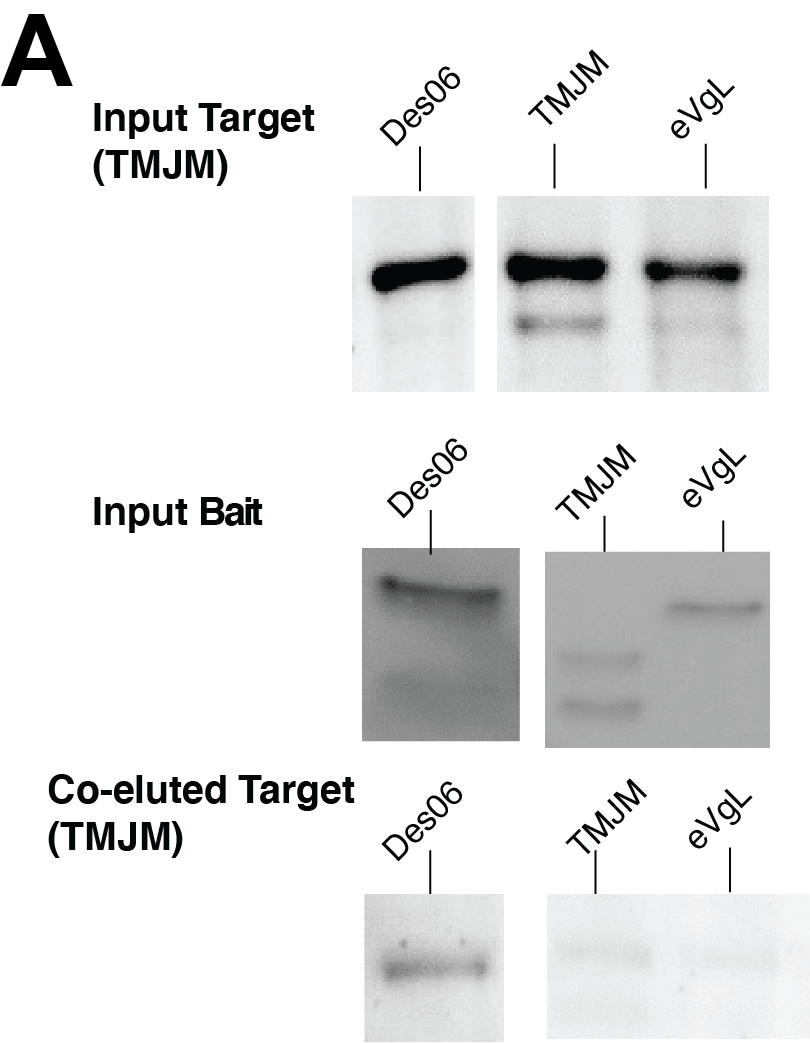

**Supplemental Figure 6. Design 6 preferentially binds TMJM in detergent-solubilized co-IP assay.**

A) Co-immunoprecipitation (co-IP) of HA-tagged target constructs with Flag-tagged TMJM bait in HEK293T cells. Cells were transiently co-transfected with plasmids encoding the Flag-tagged TMJM bait and either HA-tagged Design6, TMJM, or eVgL constructs. After 24 hours of expression, cells were lysed and membrane proteins solubilized in 1% n-dodecyl-β-D-maltoside (DDM) supplemented with 0.1% cholesterol hemisuccinate (CHS). Lysates were incubated with anti-Flag resin to isolate TMJM bait and associated proteins. Western blot analysis of HA immunoreactivity in the eluted fractions revealed preferential enrichment of HA-Design6 over HA-TMJM and HA-eVgL, indicating that Design6 exhibits stronger or more stable interaction with the TMJM bait under detergent-solubilized conditions.

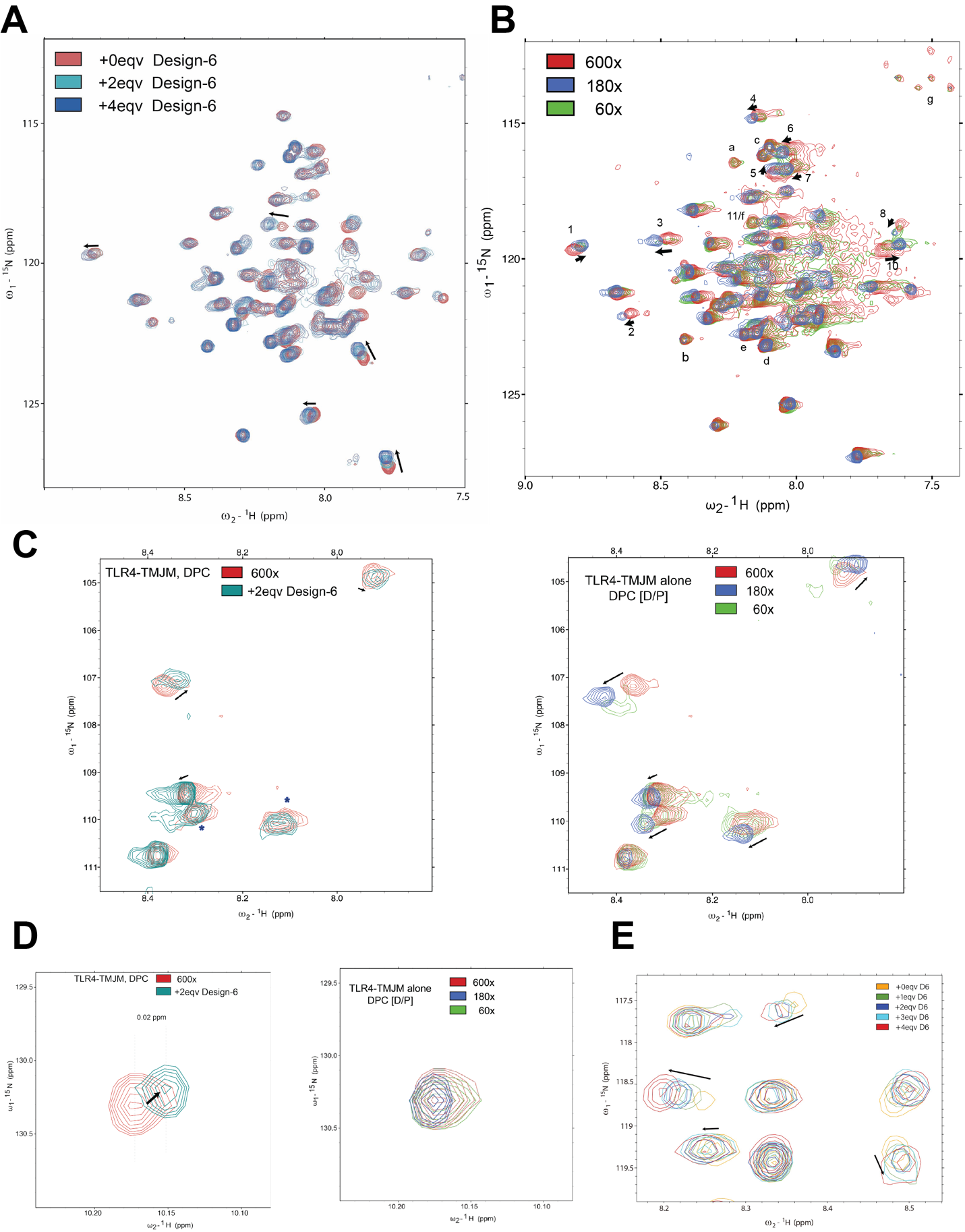

**Supplemental Figure 7. Chemical shift perturbations characterizing homotypic and heterotypic TLR4-TMJM interactions.**

A) ^1^H-^15^N HSQC spectra of 0.2 mM ^15^N His-TMJM in 60mM (300x detergent molar excess of TMJM) at 40 °C, pH 6.0, and 800 MHz alone (red) and with Design-6 peptide at 2 (aqua, 0.4mM) or 4 (blue, 0.8mM) molar equivalents. As Design-6 binder is titrated, the spectra shows a set of continuous peak shifts (marked by arrows) suggesting an interaction with TLR4-TMJM in fast-exchange.
B) The ^1^H-^15^N HSQC spectra of 0.2 mM ^15^N His-TMJM alone at 40 °C, pH 6.0, and 800 MHz at 60, 180, and 600 molar equivalents of detergent measuring detergent-dependent homo-typic interactions (12mM, 60x detergent molar excess, green; 36mM, 180x, blue; 120mM, 600x, red). Peak shifts between the most diluted sample (red) and modestly concentrated sample (blue) likely represent TLR4-TMJM’s change from a monomeric (600x) to homodimeric state (180x). The more concentrated sample (60x, green) has extensive broadening, and chemical shift perturbations not continuous with those between the 600x to 180x conditions, indicative of additional alternative states averaged from higher-order oligomerization beyond dimerization. Peaks are labeled with letters for those unperturbed by detergent titration, but shifted in TMJM’s spectra when in complex with Design-6. Peaks labelled with numbers display perturbations distinct from those induced by Design-6.
C) Glycine resonance region of spectra and chemical shift perturbation. Left, TLR4-TMJM alone at 600x DPC (red) or with 2 equivalents of Design-6 TM peptide in 36 mM ^2^H-DPC (Blue-green) from Figure 5. Right, TLR4-TMJM alone with detergent titration as in panel B.
D) Tryptophan resonances in spectra. Left, TLR4-TMJM alone at 600x DPC (red) or with 2 equivalents of Design-6 TM peptide in 36 mM ^2^H-DPC (Blue-green) from Figure 5. Right, TLR4-TMJM alone with detergent titration as in panel B.
E) Zoomed in central region of ^1^H-^15^N HSQC spectra from full titration of Design-6 peptide at 0, 1, 2, 3, and 4 molar equivalents (0mM, Yellow, 0.2mM, Green, 0.4mM Dark Blue, 0.6mM Cyan, and 0.8mM, Red) with 0.2 mM ^15^N His-TMJM in 60mM DPC (300x) including spectra in panel A.
